# From insect endosymbiont to phloem colonizer: comparative genomics unveils the lifestyle transition of phytopathogenic *Arsenophonus* strains

**DOI:** 10.1101/2024.08.06.606843

**Authors:** Mathieu Mahillon, Christophe Debonneville, Raphaël Groux, David Roquis, Justine Brodard, Franco Faoro, Xavier Foissac, Olivier Schumpp, Jessica Dittmer

## Abstract

Bacteria infecting the plant phloem represent a growing threat worldwide. While these organisms often resist *in vitro* culture, they multiply both in plant sieve elements and hemipteran vectors. Such cross-kingdom parasitic lifestyle has emerged in diverse taxa via distinct ecological routes. In the genus *Arsenophonus*, the phloem pathogens ‘*Candidatus* Arsenophonus phytopathogenicus’ (Ap) and ‘*Ca.* Phlomobacter fragariae’ (Pf) have evolved from insect endosymbionts, but the genetic mechanisms underlying this transition have not been explored. To fill this gap, we obtained the genomes of both strains from insect host metagenomes. The resulting assemblies are highly similar in size and functional repertoire, rich in viral sequences, and closely resemble the genomes of several facultative endosymbiotic *Arsenophonus* strains of sap-sucking hemipterans. However, a phylogenomic analysis demonstrated distinct origins, as Ap belongs to the “*Triatominarum*” clade whereas Pf represents a distant species. We identified a set of orthologs encoded only by Ap and Pf in the genus, including hydrolytic enzymes likely targeting plant substrates. In particular, both bacteria encode plant cell-wall degrading enzymes and cysteine peptidases related to xylellain, a papain-like peptidase from *Xylella fastidiosa*, for which close homologs are found in diverse proteobacteria infecting the plant vasculature. *In silico* predictions and expression analyses further support a role during phloem colonization for several of the shared orthologs. We conclude that the double emergence of phytopathogenicity in *Arsenophonus* may have been mediated by a few horizontal gene transfer events, involving genes first acquired from other proteobacteria including phytopathogens.

**Importance:** We investigate the genetic mechanisms of a transition in bacterial lifestyle. We focus on two phloem pathogens belonging to the genus *Arsenophonus*: *Ca.* Arsenophonus phytopathogenicus and *Ca.* Phlomobacter fragariae. Both bacteria cause economically significant pathologies, and they have likely emerged among facultative insect endosymbionts. Our genomic analyses show that both strains are highly similar to other strains of the genus associated with sap-sucking hemipterans, indicative of a recent lifestyle shift. Importantly, although the phytopathogenic *Arsenophonus* strains belong to distant clades, they share a small set of orthologs unique in the genus pangenome. We provide evidence that several of these genes produce hydrolytic enzymes that are secreted and target plant substrates. The acquisition and exchange of these genes may thus have played a pivotal role in the lifestyle transition of the phytopathogenic *Arsenophonus* strains..

## Introduction

In vascular plants, the phloem ensures the translocation of photosynthates from source to sink tissues. As a sugar-rich environment, the phloem also represents a niche for pathogenic microorganisms (1,2). In particular, bacterial infections restricted to the sieve elements (SEs, *i.e.* the phloem conductive cells) are difficult to control and cause damaging diseases worldwide (3). The causal agents are generally non-culturable and “vector-borne” as they have evolved a biphasic lifestyle, alternating between plant SEs and sap-sucking hemipteran vectors (4). Notorious examples of these bacteria include *Candidatus* (*Ca.*) species of the genera *Phytoplasma* and *Liberibacter*. This lifestyle is shared by bacteria from distant taxa, and it can emerge through distinct evolutionary routes (3,5). In the phylum *Pseudomonadota* (formerly proteobacteria), phloem pathogens have emerged either among plant-associated bacteria as in the genera *Liberibacter* and *Serratia* (1,6,7), or among insect endosymbionts as proposed for the genera *Rickettsia* and *Arsenophonus* (5,7,8). In the latter, two phytopathogens are identified, namely ‘*Ca.* Arsenophonus phytopathogenicus’ (Ap) and ‘*Ca.* Phlomobacter fragariae’ (Pf) (9–11).

Ap is transmitted by the planthoppers *Pentastiridius leporinus* and *Cixius wagneri*, and it also colonizes the SEs of sugar beet, thereby inducing the syndrome “basses-richesses” (SBR). SBR was first described in France and now occurs in Germany and Switzerland as well (12–14). Pf is transmitted by *C. wagneri* and causes strawberry marginal chlorosis (SMC) (15), first reported in France (16) and then in Japan (17). SMC was also documented in Italy, where it was associated with both Pf and Ap (18,19). Recently, Ap has been detected in potato and onion plants in Germany (20–22), as well as in sugar beets across Central Europe (23), indicating wide distribution and host range, and warning of potential novel epidemics.

While Ap and Pf are phytopathogens, other *Arsenophonus* strains are known as arthropod endosymbionts. *Arsenophonus* belongs to the family *Morganellaceae* in the order *Enterobacterales* (24). It represents one of the most widespread insect endosymbiotic genera (25), and its host range also comprises other arthropod groups such as arachnids (26). Host-symbiont interactions, tissue tropisms and transmission routes of *Arsenophonus* strains are diverse (27,28). Infections can be associated with reproductive parasitism (29), insecticide resistance (30), feeding behavior alteration (31,32) and nutritional symbiosis (33). These bacteria can multiply extra- or intracellularly, and some are maternally inherited. The best-studied species is *Arsenophonus nasoniae*, an endosymbiont of parasitoid wasps sometimes associated with “male-killing” (34). Transmission of *A. nasoniae* can be vertical, but also horizontal through multi-parasitism events (35). Horizontal transmission has also been documented for the bee endosymbiont *Arsenophonus apicola* (36). Unlike these two species, many members of the genus can only be grown in host cells or even completely resist *in vitro* culture (5). Some strains have become obligate “primary” (P) endosymbionts (37,38). This great lifestyle diversity is mirrored by various genomic architectures. While the complex genome of *A. nasoniae* consists of one large circular chromosome (> 3.5 Mb) and 7-20 extrachromosomal elements (39,40), the genomes of P-endosymbionts are highly eroded (< 1.2 Mb), lack extrachromosomal elements and display reduced metabolisms streamlined for the production of nutrients lacking from the host’s diet (37,38).

Ap and Pf exhibit traits typical for insect endosymbionts (3,5,11,41,42), which supports the idea of an ancestral insect-restricted lifestyle and the recent gain of phytopathogenicity (5). This is corroborated by phylogenetic analyses evidencing close relationships with other *Arsenophonus* strains (11). Accordingly, Ap and Pf represent ideal models to investigate the genetic changes involved in the transition from insect endosymbiont to vector-borne phloem pathogen. Probable mechanisms underlying this transition would be the acquisition of new functions, such as virulence factors, through horizontal gene transfer (HGT). To investigate this possibility, we compiled the first genomic assemblies for Pf and Ap. As both bacteria are currently non-culturable, we produced hybrid assemblies for strains of Pf (Pf-FR) and Ap (Ap-CH) using metagenomes of insects recently collected in France and Switzerland, respectively. In addition, the assembly of a second strain of Ap (Ap-FR) was obtained using short reads generated from insects sampled during an early outbreak in France. Following a genus-wide comparative analyses, we identified orthologs exclusively shared by the phytopathogenic strains that may have enabled their lifestyle transition.

## Results

### General features of the assemblies

The sizes of the genomes of Ap and Pf (2.6-2.9 Mb) are much larger than those reported for other vector-borne phloem pathogens such as phytoplasmas, liberibacters and spiroplasmas (< 1.7 Mb (43)). The main features of the three assemblies are summarized in Fig. 1A. The hybrid approach produced 50 and 33 scaffolds for Pf-FR and Ap-CH, respectively. The best assembly was obtained for the latter, as 28% of the genome is contained in the longest scaffold (806 kb) and the genome N50 is 384 kb (97 kb for Pf-FR). The genome of Ap-FR is shorter and more discontinuous with 67 scaffolds and N50 of 106 kb. The three assemblies exhibit a GC-content of c. 37%, which is similar to other facultative endosymbiotic *Arsenophonus* strains, whereas the genomes of P-endosymbionts are richer in AT (Fig. 1B). The sizes, numbers of predicted proteins (2100–2500) and pseudogenization rates (15.7-17.8 %) of the assemblies occupy an intermediate position within the genus, and align with several strains of the *Triatominarum* clade (Fig. 1B and (44)).

**Figure 1.**
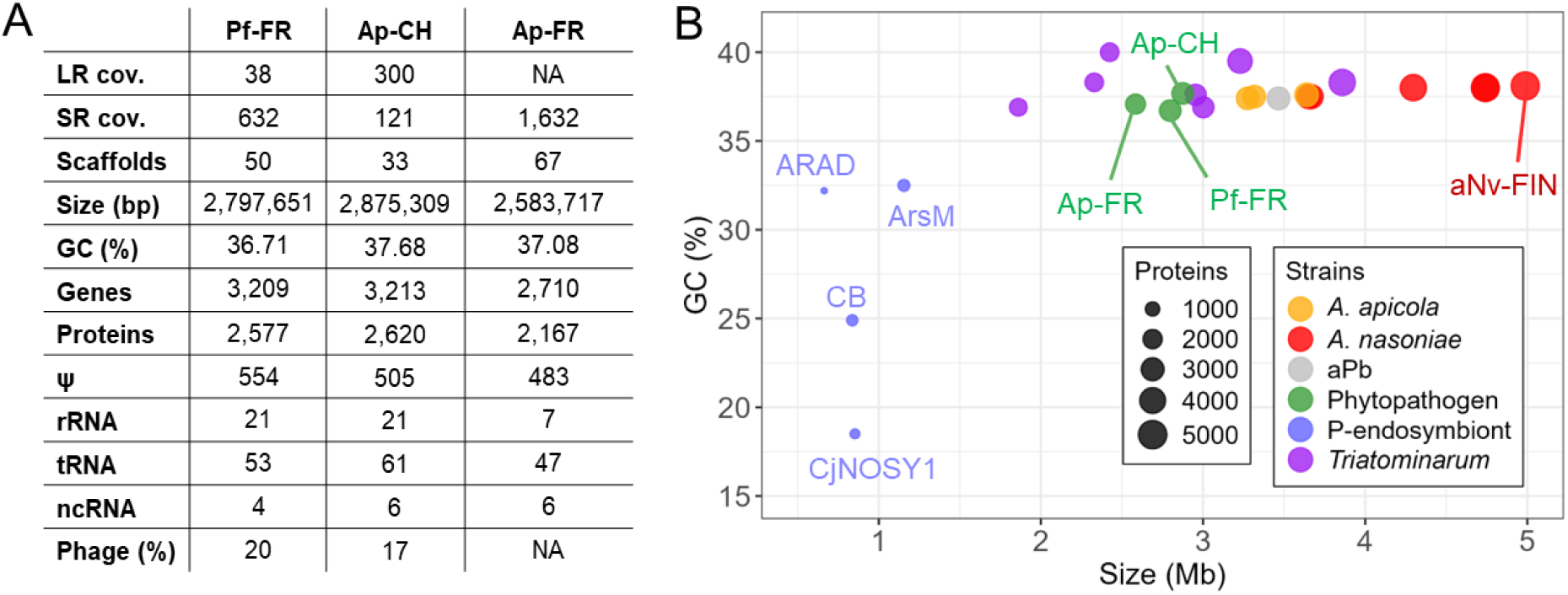
**A**: Global genomic features of Pf-FR and Ap-CH/FR. LR: long reads; SR: short reads. **B:** Distribution of GC content (in percent) and size (in megabases, Mb) for the *Arsenophonus* genomes. Each point corresponds to a single genome.

CheckM analyses confirmed a high level of genome completeness for the three assemblies: 96.76% for Ap-FR, 97.84% for Pf-FR and 99.46% for Ap-CH. In the latter, seven complete rRNA operons were identified, while a previous study found only six copies of the 16S rRNA (45). Three complete rRNA operons were detected for Pf-FR, but this genome likely contains at least seven rRNA operons as additional rRNA genes occur at scaffold ends. Both Ap genomes show extensive similarities (Fig. S1), hence some of the finer-scale analyses presented herein will focus only on Ap-CH.

### Phage and plasmid regions

Despite high sequencing coverages and multiple assembly approaches, the three assemblies could not be closed, consistent with previous *Arsenophonus* genome projects necessitating DNA from pure cultures (40,46). The assemblies of Ap and Pf may thus contain extrachromosomal elements, as reported for other strains of the genus (Table 1 and (44)). Several *Arsenophonus* extrachromosomal elements were recently characterized as “phage-plasmid”, harboring both phage and plasmid modules (40,47), and some probably correspond to helper prophages and their associated satellites (48).

**Table 1.** Genomes of *Arsenophonus* strains used in this study.

| Strain | Host species | Size (Mb) | Status * | P ** | Accession | Ref |
| --- | --- | --- | --- | --- | --- | --- |
| **Facultative endosymbionts** |  |  |  |  |  |  |
| *Ca. A. phytopathogenicus* Ap-CH | *Pentastiridius leporinus* | 2.9 | S | NA | JBDPZB000000000 | This study |
| *Ca. A. phytopathogenicus* Ap-FR | *Pentastiridius leporinus* | 2.6 | S | NA | JBDPZA000000000 |  |
| *Ca. P. fragariae* Pf-FR | *Cixius wagneri* | 2.8 | S | NA | JBDPYZ000000000 |  |
| *A. nasoniae* aNv-CAN | *Nasonia vitripennis* | 5.1 | C | 20 | GCF_029873515.1 | (44) |
| *A. nasoniae* aNv-CH | *Nasonia vitripennis* | 4.7 | C | 16 | GCF_029873535.1 | (36) |
| *A. nasoniae* aNv-DSM | *Nasonia vitripennis* | 3.6 | S | NA | GCF_000429565.1 | (46) |
| *A. nasoniae* aNv-FIN | *Nasonia vitripennis* | 5.0 | C | 17 | GCF_004768525.1 | (40) |
| *A. nasoniae* aNv-UK | *Nasonia vitripennis* | 4.7 | C | 16 | GCF_029873555.1 | (44) |
| *A. nasoniae* aPv | *Pachycrepoideus vindemmiae* | 4.3 | C | 18 | GCF_029873495.1 | (44) |
| *A. nasoniae* alh | *Ixodiphagus hookeri* | 3.7 | C | 7 | GCF_029873455.1 | (44) |
| *A. apicola* aApi-US | *Apis mellifera* | 3.6 | C | 6 | GCF_020268605.1 | (50) |
| *A. apicola* aApi-CH | *Apis mellifera* | 3.3 | S | NA | GCF_903968575.1 | (36) |
| *A. apicola* aApi-AU | *Apis mellifera* | 3.2 | C | 5 | GCF_029906405.1 | (44) |
| *A. triatominarum* ATi | *Triatoma infestans* | 3.9 | S | NA | GCF_001640365.1 | - |
| *Arsenophonus* sp. aPb | *Polyommatus bellargus* | 3.5 | C | 4 | GCF_029873475.1 | (51) |
| *Ca. Arsenophonus* sp. ENCA | *Entylia carinata* | 3.2 | S | NA | GCF_002287155.1 | (114) |
| *Ca. Arsenophonus* sp. ARAF | *Aleurodicus floccissimus* | 3.0 | S | NA | GCF_900343025.1 | (38) |
| *Ca. Arsenophonus* sp. Hangzhou | *Nilaparvata lugens* | 2.9 | S | NA | GCF_000757905.1 | (115) |
| *Ca. Arsenophonus* sp. Ash | *Aphis craccivora* | 2.4 | C | 1 | GCF_013460135.1 | - |
| *Ca. Arsenophonus* sp. Asia-II-3 | *Bemisia tabaci* | 2.3 | S | NA | GCF_004118055.1 | (116) |
| *Ca. Arsenophonus* sp. MEDQ21 | *Bemisia tabaci* | 1.9 | S | NA | GCF_902713415.1 | - |
| **Primary endosymbionts** |  |  |  |  |  |  |
| *Ca. A. melophagi* ArsM | *Melophagus ovinus* | 1.1 | S | NA | NA*** | (33) |
| *Ca. Arsenophonus* sp. CjNOSY1 | *Ceratovacuna japonica* | 0.9 | C | 0 | GCA_024349725.1 | (117) |
| *Ca. A. lipoptenae* CB | *Lipoptena cervi* | 0.8 | C | 0 | GCF_001534665.1 | (37) |
| *Ca. Arsenophonus* sp. ARAD | *Aleurodicus dispersus* | 0.6 | C | 0 | GCF_900343015.1 | (38) |

\*S: scaffolds; C: closed. \*\*Plasmid number. \*\*\*Genome available online: http://users.prf.jcu.cz/novake01

Four regions in Ap-CH and one in Pf-FR exhibit synteny with sequences from the *Arsenophonus* extrachromosomal elements (Fig 2A, black blocks, and Table S2). Plasmid genes are present both inside and outside these regions in both assemblies (Table S3). Phaster predicted 18 and 26 phage regions for Ap-CH and Pf-FR, respectively, representing 17-20% of the genomes (Fig. 2A, green, blue and red blocks). These repetitive regions likely contributed to the assembly breaks, as indicated by their locations at scaffold ends (Fig. 2A, inner ribbons). The majority of these regions are short and predicted as “incomplete”, harboring only a few viral genes (Fig. 2B). Their status should thus be taken with caution since they may represent phage relics or partial sequences. Conversely, eight regions for each strain are classified as “questionable” and even “intact”, and could represent functional phages.

**Figure 2:**
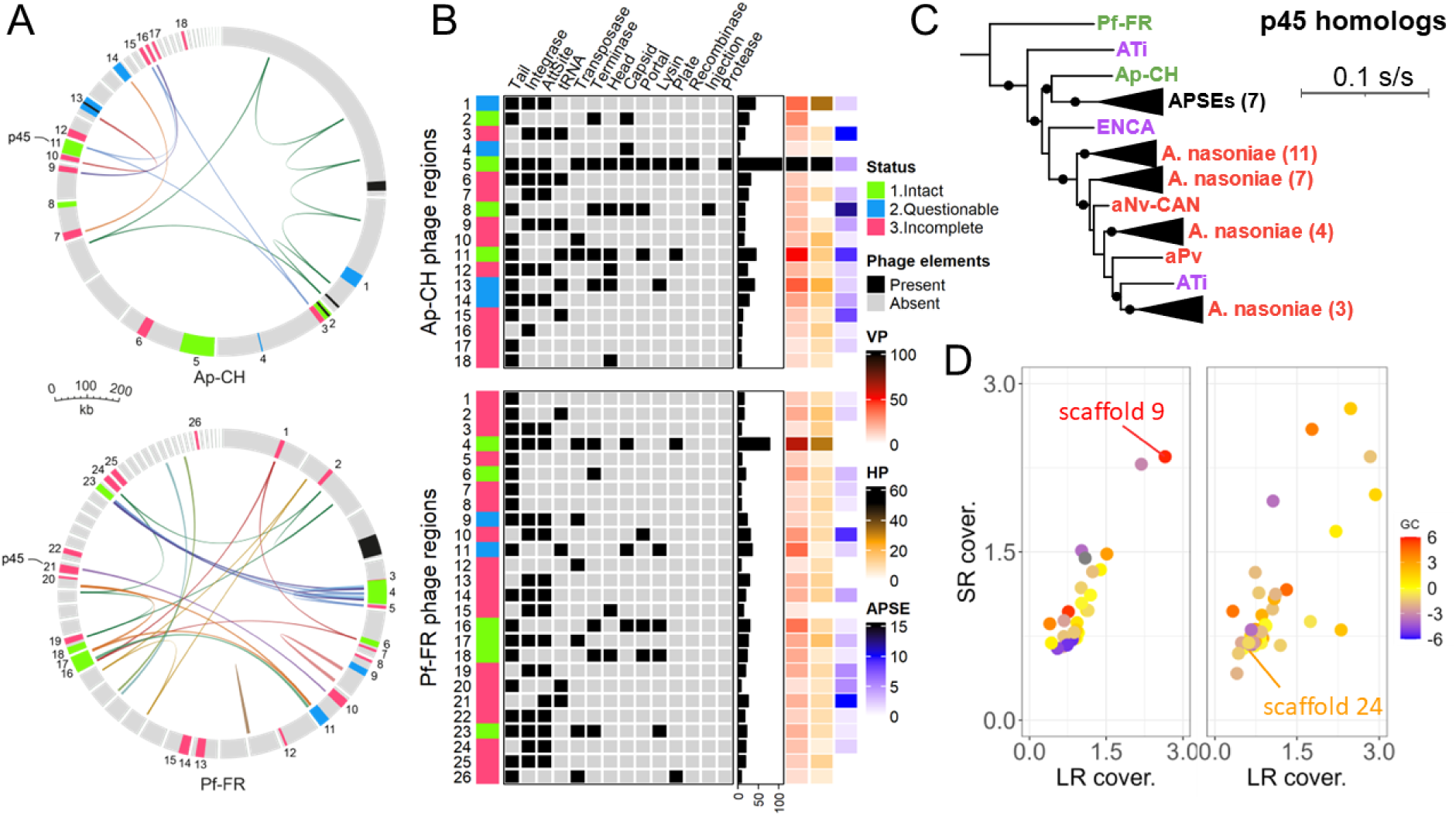
Viral content in Ap-CH and Pf-FR. **A:** Schematic representation of the genome assemblies of Ap-CH (top) and Pf-FR (bottom). Each assembly is represented as a circle accommodating the scaffolds. Black blocks indicate regions syntenic to *Arsenophonus* extrachromosomal elements. Other colored blocks represent phage regions (> 1 kb) identified by Phaster. Inner ribbons connect syntenic blocks (> 5 kb). **B:** Composition of the phage regions. VP: number of viral proteins; HP: number hypothetical non-viral proteins; APSE: number of APSE proteins. The barplot represent the size of each region in kb. **C:** Maximum-likelihood phylogenetic tree of the p45 homologs, built with the model JTT+F+I+G4. Black circles on branches indicate >50% bootstrap support. The scale is given in substitutions per site. Polymerases from *Citrobacter koseri* (WP_200053203.1) and *Salmonella enterica* (EFS9905187.1) were used to root the initial tree. **D:** Normalized long-read (LR) and short-read (SR) coverages for the scaffolds of Ap-CH (left) and Pf-FR (right). The color scale indicates the difference in GC-content from the assembly average.

Importantly, one intact phage DNA polymerase was detected in both hybrid assemblies (ApCH_3206 and Pf-FR_1270). They represent homologs of p45 (> 87% aa id.), the polymerase of *Acyrthosiphon pisum* Secondary Endosymbiont (APSE) phages (49). Similar homologs are found in several other *Arsenophonus* strains (Fig. 2C). Entire APSE modules have been previously identified in *Arsenophonus* genomes, in which they form mosaics with other phage modules (49). Likewise, in Ap-CH and Pf-FR, numerous APSE genes are found throughout the assemblies (Fig. 2B, column “APSE”). In particular, the p45 homologs are located within gene clusters syntenic to APSE replicative modules 1 and 2 (Fig. S2). In the case of Ap-CH, these modules are part of the intact phage region 11 on scaffold 9, which also contains the incomplete phage region 10 and several plasmid genes (Table S3). This scaffold exhibits a high GC-content of 43% and a high sequencing coverage (Fig. 2D), indicating that it may be a multi-copy phage-plasmid element. In Pf-FR, the APSE replicative modules are located within the incomplete phage region 21 on scaffold 24, which shows no discrepancy in coverage and GC-content compared to other scaffolds (Fig. 2D), and might hence represent a single copy region. These modules could be associated with an integrated prophage since the rest of the scaffold consists of non-viral genes.

Unfortunately, the APSE replicative modules of both assemblies are located at scaffold ends (Fig. S2), limiting further characterization.

### Phylogenomic relationships

A phylogenomics analysis combining Ap, Pf and other facultative symbionts revealed two basal clades within the genus (“I” and “II”, Fig. 3), consistent with previous studies (50,51). An analysis including the P-endosymbionts resulted in a similar tree topology, albeit with reduced confidence due to long-branch attraction caused by the reduced genomes (Fig. S2). Clade I accommodates the hymenoptera-infecting species *Nasoniae*, *Apicola* and the butterfly-infecting strain aPb, whereas clade II, or “*Triatominarum*” clade (44), comprises hemipteran endosymbionts.

**Figure 3.**
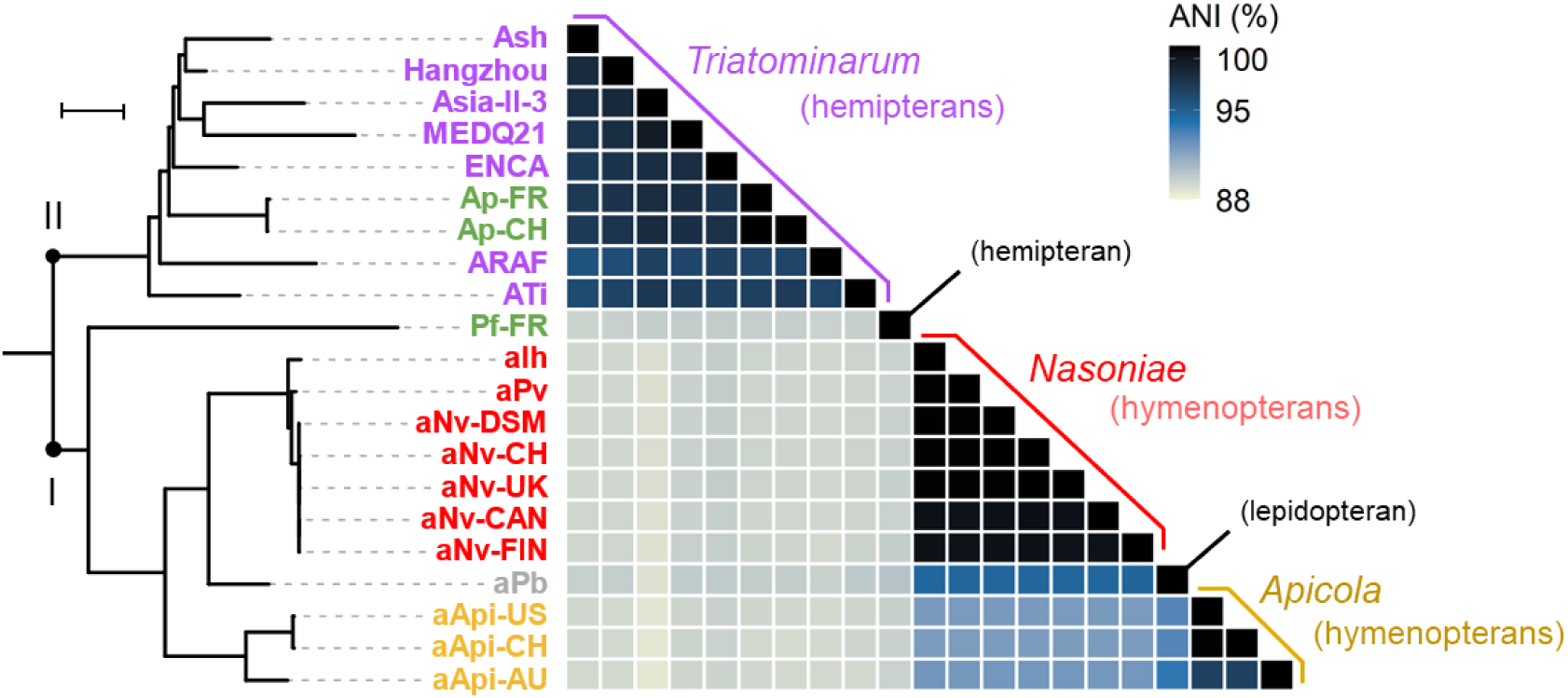
Phylogenomic and taxonomic analyses of facultative endosymbiotic *Arsenophonus* strains. **Left:** ML phylogenomic tree based on 118 single copy protein-coding genes. All branches have > 90% bootstrap support. Genes from *Providencia stuartii* and *Proteus mirabilis* were used to root the initial tree. The tree scale represents 0.01 substitution per site. Basal clades are indicated by latin numbers. **Right:** Matrix of pairwise average nucleotide identities (ANI, in percent). The host ranges are given in parenthesis.

In the tree, Ap and Pf unambiguously represent distinct species. The Ap strains are deeply embedded within the *Triatominarum* clade (bootstrap support: 100%), which members can be considered as strains of the same “species complex” since they share very high ANI levels (> 95% (52)). In contrast, Pf is an early-branching member of clade I (bootstrap support: 98%) and shares low ANI levels (< 91%) with the other strains, indicating that it represents a distinct species within the genus.

### Basal biosynthetic capacities

The diversity of lifestyles and genome sizes among *Arsenophonus* strains is reflected by substantial variations in basal biosynthetic capacities (Fig. 4). There is a clear erosion gradient from the well-furnished genomes of clade I culturable strains to the reduced genomes of P-endosymbionts. Notably, the strain Hangzhou from the *Triatominarum* clade exhibits a basal biosynthetic repertoire almost equivalent to that of clade I strains, consistent with the idea that the genus ancestor was a free-living insect-associated bacterium (36). Ap and Pf have experienced an intermediate level of erosion and have lost many biosynthetic pathways. Nevertheless, these strains have retained more capacities than most members of the *Triatominarum* clade, showing a repertoire highly similar to the strain ENCA.

**Figure 4.**
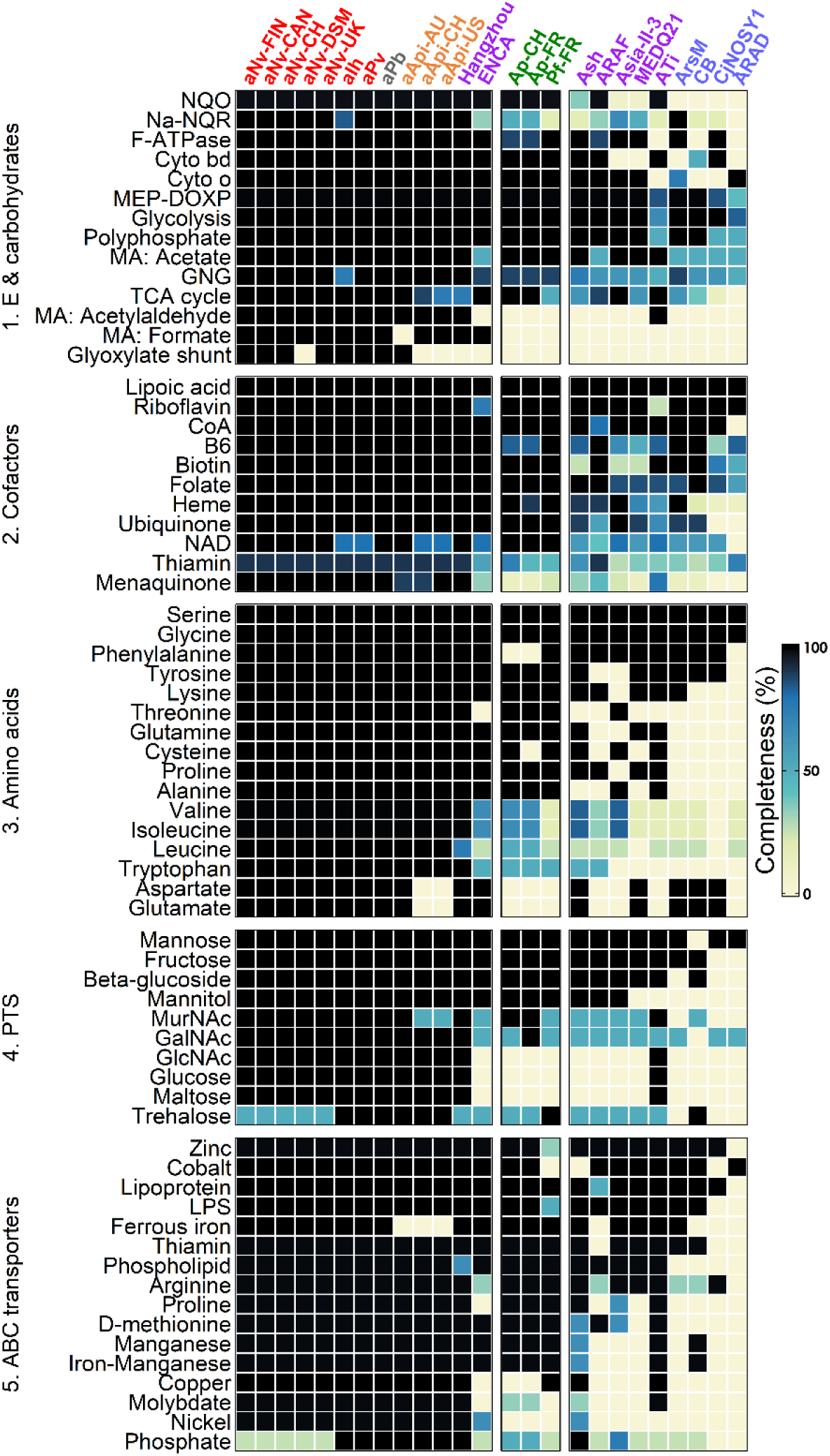
Biosynthetic capacities and transmembrane transport systems among *Arsenophonus* strains. Rows indicate the pathway products or the transported substrates, while columns indicate the strains. Colors indicate the completeness according to the KEGG DB. Only pathways complete in at least one strain are shown. B6: vitamin B6; Coa : coenzyme A; Cyto bd: cytochrome bd complex; Cyto o : cytochrome o ubiquinol oxidase; NaNQR: Na^+^-NADH-ubiquinone oxidoreductase; NAD: nicotinamide adenine dinucleotide; NQO: NADH quinone oxidoreductase; MA: mixed acid fermentations; GNG: Gluconeogenesis GlcNAc; N-acetylglucosamine. GalNAc : N-acetylgalactosamine. N-acetylmuramic acid: MurNAc; LPS: lipopolysaccharide. MEX-DOXP: non-mevalonate pathway.

In terms of energy transfer, Ap and Pf have intact NADH quinone oxidoreductase and cytochromes, but the F-type ATPase is incomplete in the Ap strains. For the carbohydrate metabolism, both Ap and Pf can rely on the glycolysis and non-mevalonate pathway, and the Ap strains also have a complete tricarboxylic acid cycle. Interestingly, the phytopathogenic strains have retained the capacity to ferment sugars into acetate, while they have lost other mixed acid pathways. Ap and Pf can produce most cofactors, and this capacity could be involved in nutritional symbiosis with their cixiid hosts feeding on nutrient-limited sap (38,53). On the other hand, the phytopathogenic strains have lost the ability to produce numerous amino acids, indicating that these compounds must be obtained from the hosts. Ap and Pf encode several phosphotransferase systems, many ABC transporters and a siderophore (Fig. 4 and Table S4). This capacity for nutrient acquisition could create phloem imbalances and partially explain the plant symptoms associated with SBR and SMC, as suggested for other phloem-infecting bacteria (54–56).

### Genes for virulence and symbiosis

Previous studies have evidenced a complex genetic arsenal dedicated to the virulence and symbiotic lifestyle of *Arsenophonus* strains in their insect hosts (44,46). Similar to basal capacities, a gradient in this gene arsenal is observed among the strains (Fig. 5). As facultative endosymbionts, Ap and Pf have retained a large portion of this arsenal, which is likely required during the infection of cixiids in order to survive in the gut, cross cellular barriers, evade immunity and reach the eggs and salivary glands (46).

**Figure 5.**
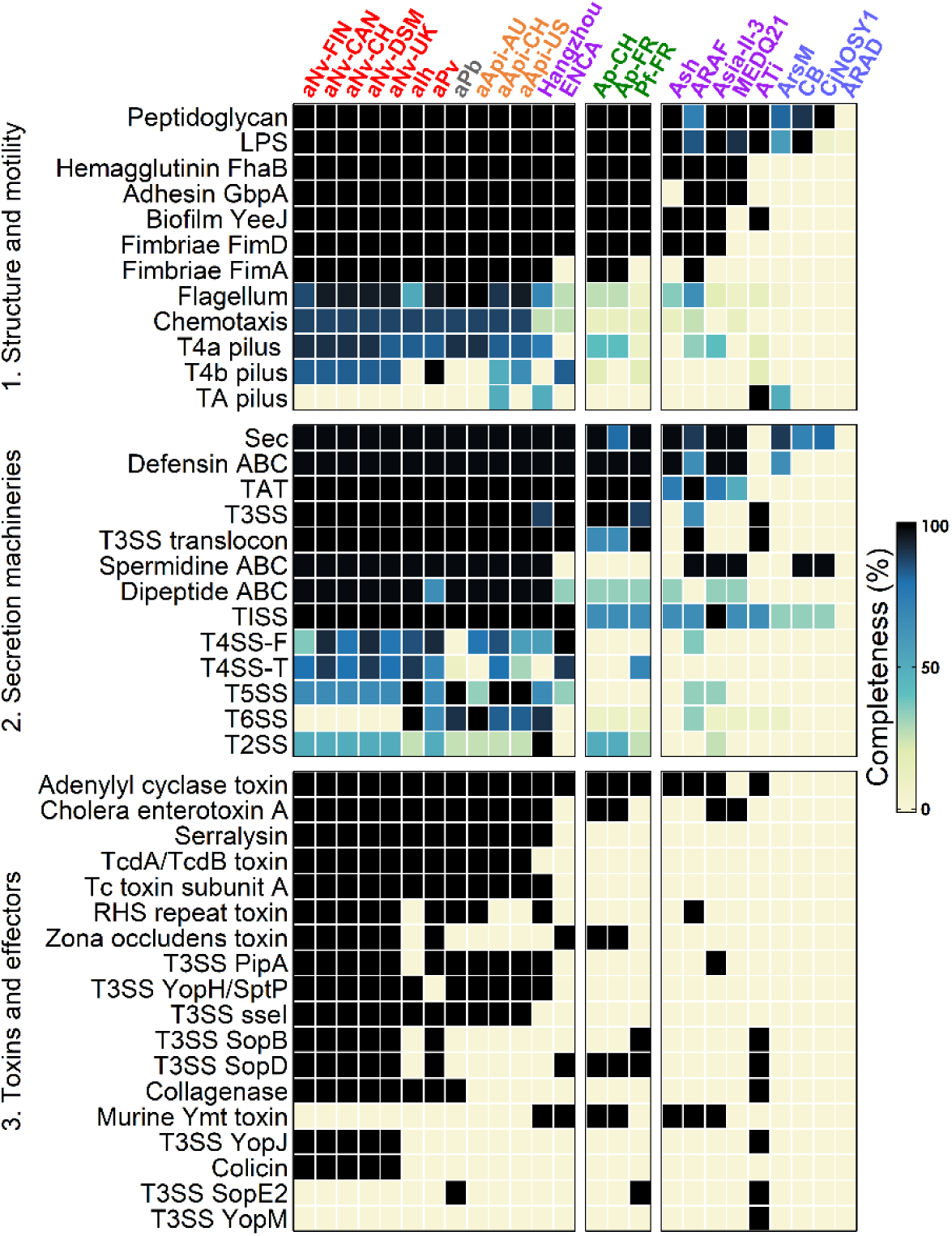
Gene arsenal involved in virulence/symbiosis in the *Arsenophonus* strains. Rows indicate the pathway/protein products, and columns represent the strains. Colors indicate the pathway completeness according to the KEGG DB. Only pathways complete or near-complete in at least one strain are shown. TA: tight adherence; TAT: twin arginine translocation; Sec: general secretion; SS: secretion system.

Ap and Pf possess the genetic capacities to produce lipopolysaccharides and peptidoglycan, consistent with studies reporting typical gram-negative cell walls (41,57,58). Both strains also harbor genes involved in adhesion but lack the capacity for flagellum, pilus and chemotaxis, which may explain their fastidious *in planta* multiplication (14,58). The pathways for general secretion (SEC) and twin arginine translocation (TAT) are both conserved in Ap and Pf. These bacteria also have a defensin ABC transporter, and the type 3 secretion system (T3SS) is complete for both Ap strains and almost complete for Pf-FR (lacking only *sctQ*). Conversely, other secretion systems appear absent or incomplete. A few toxins and effectors are predicted in Ap and Pf genomes, and those probably play a role during the insect stage given their presence in non-phytopathogenic strains.

### Genes specific to Ap and Pf

The molecular mechanisms underlying the phytopathogenicity of Ap and Pf have not been studied, and it is unclear whether similar or distinct invasion strategies are used. Nevertheless, considering the unique lifestyle of Ap and Pf in the genus, it is imaginable that the genes involved in phloem colonization are lacking in other *Arsenophonus* strains. To identify such “phytopathogen-specific” genes, an orthology clustering analysis was first conducted, resulting in 93% the *Arsenophonus* pangenome clustered into 4,947 orthogroups (OGs) (Fig. 6A). The core-genome is relatively small, comprising only 3,337 genes in 119 OGs. Importantly, with the exception of ATi and to a lesser extent ENCA, the percentages of strain-specific genes (*i.e.* unassigned genes and genes in strain-specific OGs) are quite low in all strains (Fig. 6B). Only 159 genes are specific to Ap and/or Pf (Fig. 6C). Of these, some represent regulatory and mobile genetic elements, but the majority encode short hypothetical proteins (HPs) with no known homolog or distant homologs in diverse enterobacteria (Table S5).

**Figure 6.**
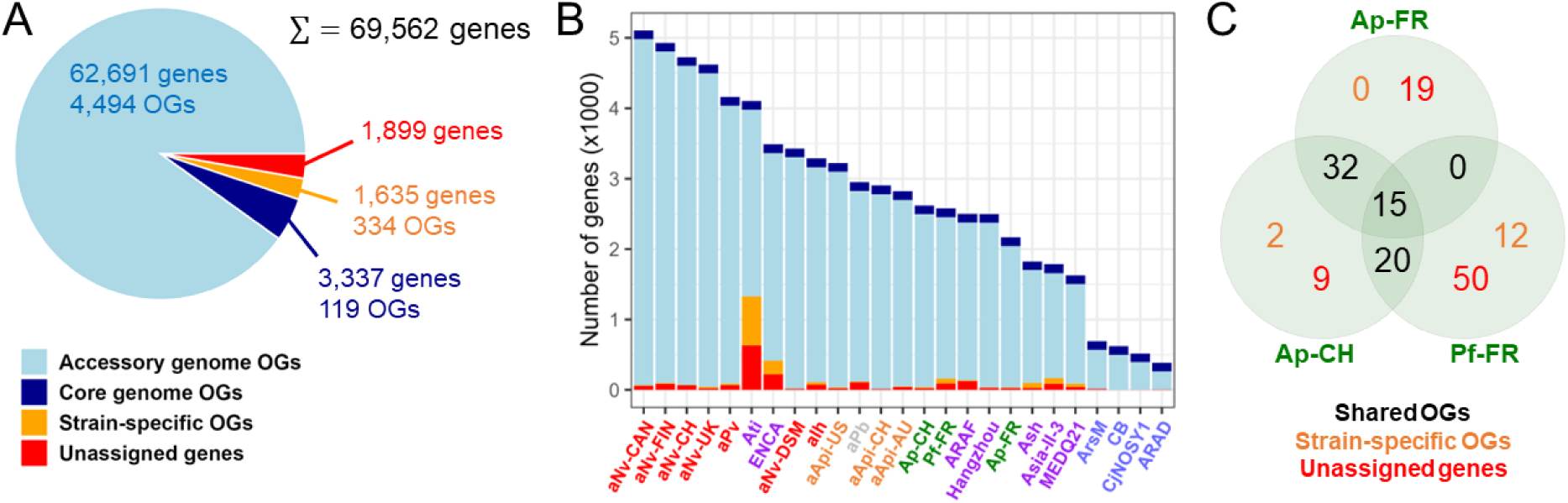
Identification of genes specific to Ap and Pf in the genus. **A**: Orthology clustering of the *Arsenophonus* pangenome. **B:** Distribution of genes in each *Arsenophonus* strains. **C:** Venn diagram of genes only found in Ap and Pf.

Importantly, 35 genes in eight OGs are exclusively shared by Pf-FR and Ap-CH/FR in the genus (Fig. 6C and Table 2). Proteins from **OG3381** are short Zn metallopeptidases with no known homolog. **OG2682** corresponds to phage lysis proteins with distant homologs in diverse enterobacteria (Fig. 7A). **OG3842** are homologs of the transcriptional repressor *DicA*, with the closest homologs found in *Enterobacter*, the endophyte *Kosakonia pseudosacchari* and the two phytopathogens *Erwinia billingiae* and *Pectobacterium brasiliense* (Fig. 7B). Likewise, **OG4231** are putative chaperones with homologs found in both phytopathogenic (*Erwinia tracheiphila and Pantoea stewartii*) and non-phytopathogenic enterobacteria (Fig. 7C). Proteins from **OG3860** contain the Domain of Unknown Function 3757 (DUF3757) with distant homologs in the phytopathogenic genera *Ralstonia*, *Pseudomonas*, *Lonsdalea* and *Burkholderia* (Fig. 7D). Similar homologs were also identified among unassigned genes (ApCH_1314 and PfFR_1763, Table S5).

**Figure 7.**
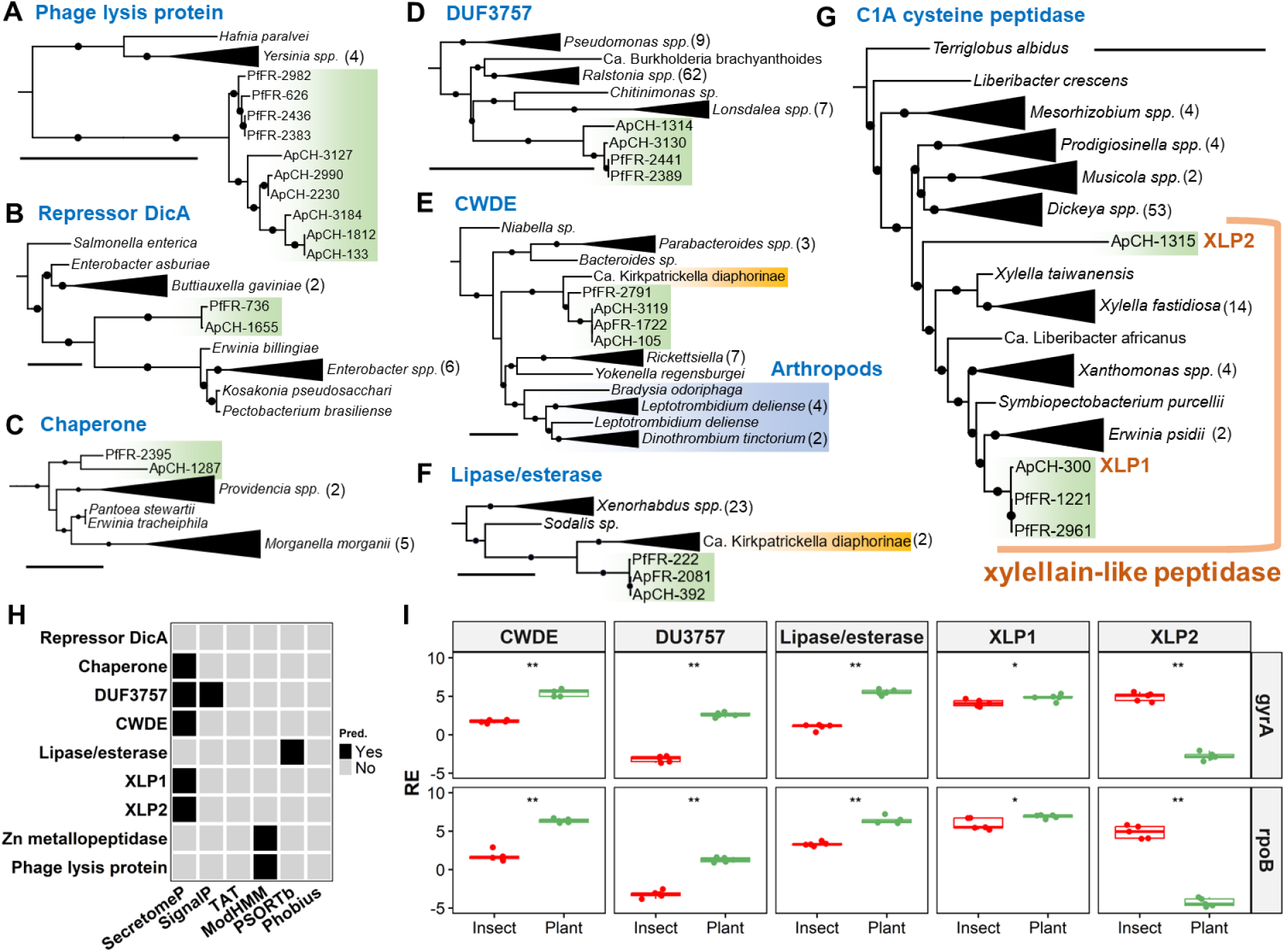
Characterization of a subset of OGs specific to Ap and Pf in the *Arsenophonus* pangenome. A-G: Maximum-likelihood phylogenetic trees. Only the closest homologs are shown in the trees. Black circles on branch indicate >50% bootstrap support. The black scales represent one substitution per site. Numbers in brackets indicate the number of genes present in the collapsed clades. *Arsenophonus* genes are highlighted in green. Arthropod genes are highlighted in blue. Genes from *Ca.* K. diaphorinae are highlighted in yellow. The trees were built using the models WAG+I+G4, JTT+I+G4, JTT+G4, VT+I+G4, LG+F+I+G4, VT+I+G4 and WAG+I+G4, respectively. H: *In silico* predictions for secretion and subcellular localization signals. I: Box plots of relative expression (in log2) for selected genes of Ap-CH in insect (red) and plant (green) tissues. Expression levels (n = 5) are given relative to the expression of *gyrA* (top) or *rpoB* (bottom). Single and double asterisks indicate p-values < 0.05 and <0.01, respectively, based on Wilcoxon Rank Sum tests.

**Table 2.** Description of the OGs shared exclusively by Ap and Pf among *Arsenophonus* strains. Blastp results are shown for the longest gene of each OG.

| OG | Size (aa) | Domain (E-value) | Annotation | Best BlastP hit |  |  |  |
| --- | --- | --- | --- | --- | --- | --- | --- |
|  |  |  |  | Accession | Size (aa) | ID (%) | Species/Strain |
| 2682 | 150 | PF03245 (6e-27) | Phage Rz lysis protein | WP_249540969.1 | 154 | 41.2 | *Escherichia coli* E15 |
| 3381 | 109 | COG2321 (2e-3) | Zn metallopeptidase | No hit |  |  |  |
| 3615 | 533 | PF14587 (1.2e-11) | O-glycosyl hydrolase | WP_319806414.1 | 532 | 86.5 | *Ca. K. diaphorinae* |
| 3842 | 132 | PRK09706 (6e-8) | Repressor DicA | WP_265910051.1 | 240 | 60.0 | *Erwinia billingiae* LS-1 |
| 3860 | 142 | DUF3757 (3e-17) | Unknown | CBJ39888.1 | 140 | 53.0 | *Ralstonia solanacearum* CMR15 |
| 3863 | 271 | COG4870 (2e-45) | C1A cysteine peptidase | WP_233592545.1 | 271 | 76.7 | *Erwinia psidii* LPF 681 |
| 3866 | 504 | PF09994 (3e-18) | Alpha/beta hydrolase | WP_319806123.1 | 590 | 40.4 | *Ca. K. diaphorinae* |
| 4231 | 99 | COG2833 (9.2e-1) | Molecular chaperone | WP_233480985.1 | 96 | 73.7 | *Erwinia tracheiphila* MDcuke |

Curiously, **OG3615, OG3866** and two unassigned genes from Pf show homology to genes from the psyllid endosymbiont *Ca.* Kirkpatrickella diaphorinae (59). **OG3615** encodes O-glycosyl hydrolases of the CAZyme family GH30. The closest homologs are found in the poorly-characterized subfamily GH30-6 (60), with the sole characterized member belonging to *Parabacteroides gordonii* (WP_028726386.1). This enzyme is active on pNP-β-D-cellobioside (61), indicating that OG3615 could represent cell wall degrading enzymes (CWDEs) targeting plant polysaccharides. Intriguingly, homologs are encoded by arthropod endosymbionts (*Yokenella* and *Rickettsiella*), gut commensals (*Parabacteroides* and *Bacteroides*) but also a species of fungus gnat (*Bradysia odoriphaga*) and several mites of the genera *Leptotrombidium* and *Dinothrombium* (Fig. 7E). **OG3866** represents hydrolases of unknown function distantly related to proteins from diverse non-phytopathogenic enterobacteria (Fig. 7F). An unassigned gene from Pf-FR (PfFR_3139) is also closely related to this OG. These hydrolases putatively represent lipases/esterases as they carry the motif “GxSxG” (62).

Strikingly, **OG3863** represents a set of C1A cysteine peptidases closely related to xylellain (AE003869, 65% aa id.), a papain-like peptidase described in *Xylella fastidiosa* (63). Hence, OG3863 and related proteins are hereafter dubbed “xylellain-like peptidases” (XLPs). A second, distinct XLP is found among unassigned genes of Ap-CH (ApCH_1315, “XLP2”) and two additional XLP pseudogenes are present in Pf-FR (PfFR_663 and 664). Importantly, XLPs are shared among distantly-related phytopathogenic proteobacteria: *Erwinia psidii*, *Xylella taiwanensis*, *Xanthomonas albilineans* and “pseudalbilineans” and *Ca.* Liberibacter africanus (Fig. 7G). A homolog is also encoded by the leafhopper endosymbiont *Symbiopectobacterium purcellii*. The XLPs are closely related to enzymes from the phytopathogenic genera *Dickeya* and *Musicola*, and from pectinolytic bacteria of the genus *Prodigiosinella* (64). Distant homologs are found in the endophyte *Liberibacter crescens* and nodule-forming *Mesorhizobium*.

### *In silico* predictions and gene expression analyses

Several *in silico* prediction tools were used to assess potential secretion and subcellular localization for the longest representative of each OG exclusively shared by Ap and Pf (Fig. 8A). A non-classical secretion was predicted by SecretomeP for the chaperone, CWDE, XLPs and DUF3757. SignalP detected a signal peptide for the DUF3757 product, indicating a putative Sec-dependent secretion. Transmembrane regions were evidenced by ModHMM in the Zn metallopeptidase and phage lysis protein, and PSORTb predicted a localization in the outer membrane for the lipase/esterase.

The expression levels of several OGs, in particular the hydrolytic enzymes and DUF3757, were determined by RT-qPCR analyses on RNA samples extracted from Ap-infected stems of periwinkle (*Catharantus roseus*) and female *P. leporinus* bodies. Importantly, expression profiles for all tested genes were similar regardless of the reference gene (*gyrA* or *rpoB*). Significantly higher expressions *in planta* were detected for the DUF3757, lipase/esterase and CWDE, and to a lesser extend XLP1. In contrast, XLP2 expression was much higher in insect tissues.

### Discussion

For this first genomic analysis of Ap and Pf, three assemblies were obtained from planthopper metagenomes. Despite the use of ONT sequencing, the genomes remain at the scaffold level, partly due to the presence of repetitive viral sequences. Axenic cultures of Ap and Pf will probably be required to obtain longer sequencing data and achieve gap closure, and this will be valuable to further explore the putative phage plasmids. Specifically, the APSE modules deserve more scrutiny as they seem widespread in *Arsenophonus* and are known to play important biological roles in other insect endosymbionts (65,66).

Comparison of global genomic features and biosynthetic capacities indicate that Ap and Pf are highly similar to several other *Arsenophonus* strains, especially facultative endosymbionts of sap-sucking hemipterans. Moreover, Ap and Pf harbor very low fractions of strain-specific genes, which reinforces the idea of a recent shift from an insect-strict to a dual insect-phytopathogenic lifestyle. Phylogenomic and ANI analyses further support this hypothesis at least for Ap, as evidenced by very close relationship with other strains from the *Triatominarum* clade. Importantly, Ap and Pf share a small set of eight OGs unique in the *Arsenophonus* pangenome. Of these OGs, little can be speculated about the putative chaperones, phage lysis proteins, metallopeptidases, DUF3757 and lipases/esterases, although similar proteins have been previously characterized as virulence factors in bacterial phytopathogens (67–69). In contrast, a role in the plant stage of Ap and Pf can be more confidently assumed for the CWDEs and XLPs.

The shared CWDEs belong to the large glycoside hydrolase family GH30, which include enzymes used by phytopathogens to degrade cellulose, hemicellulose and pectin (70–72). Both Ap and Pf cause physiological alterations in plants including SE obstruction (57,58), which might be targeted by the CWDEs to ease systemic invasion. These enzymes might also have nutritional functions through the degradation of plant polysaccharides into consumable forms. This is supported by the presence of homologs in other arthropod endosymbionts and gut commensals. Surprisingly, homologs of these CWDEs are also found in several mites and a fungus gnat, indicating probable cross-kingdom HGT. Notably, HGT between bacteria and arthropods has been previously documented, and include genes crucial for plant adaptation (73,74).

The occurrence of XLPs in diverse phytopathogenic proteobacteria clearly suggests a role in plant tissue colonization. Although the xylellain has been purified and its structure resolved, its substrate remains unknown (63). The XLPs represent a sister clade of enzymes found in soft rot agents, plant endophytes and rhizobia, suggesting that plant proteins represent targets of all these C1A cysteine peptidases. Importantly, all XLP-encoding phytopathogenic bacteria infect the xylem, phloem or both tissues (75–78). Therefore, these enzymes may specifically act on vasculature proteins. The XLPs are hypothetically secreted, similar to several cysteine peptidases characterized in other phytopathogens, which are T3SS effectors targeting the plant immunity (79). However, low homology between these effectors and XLPs precludes further comparison (data not shown). Intriguingly, an XLP is found in *S. purcellii* SyEd1T, a leafhopper endosymbiont with no known plant stage (80). It should be noted that the closely-related strain BEV is plant transmitted (81), and similar strains have been recently detected in potato (82), indicating that SyEd1T might also have a plant stage. Interestingly, in contrast to Ap XLP1, Ap XLP2 appear to be more expressed in insect than plant tissues. This could be associated with a specific role in the plant adaptation of the cixiid vector through the digestion of phloem proteins, as hypothesized for peptidases encoded by sap-sucking hemipterans and their endosymbionts (83–85).

It is likely that some OGs found exclusively in Ap and Pf were transferred from one strain to the other by direct HGT, which might have taken place in a co-infected *C. wagneri* specimen as this planthopper can host both phytopathogens. Prior to these exchanges, some shared OGs were likely acquired from other proteobacteria including phytopathogens with no known insect host (e.g. *Erwinia*). Importantly, Ap and Pf might have developed the ability to survive in the phloem before acquiring phytopathogenic traits. Indeed, diverse non-phytopathogenic endosymbionts use SEs for horizontal transmission between hemipterans (86–89), and such a plant-mediated transmission would explain the diversity of sap-sucking hosts harboring closely-related strains of the *Triatominarum* clade. From this capacity to survive in the phloem, Ap and/or Pf might have been in contact with and acquired genes (e.g. XLP) from phytopathogens. Noteworthily, several gene may have been exchanged with *Ca.* K. diaphorinae, an endosymbiont of the *Liberibacter* vector *Diaphorina citri*. HGT with this bacterium could have occurred in a co-infected hemipteran. HGT *in planta* is also imaginable, as *D. citri* is closely related to *Asaia*, a genus of arthropod endosymbionts horizontally transmitted via plants (90,91).

In conclusion, our results strengthen the idea of an “insect first” scenario during the evolution of Ap and Pf towards becoming vector-borne phytopathogen. A limited number of HGT events probably acted as key mechanisms in this lifestyle transition, which aligns with recent studies associating HGT with the shift from non-vascular to vascular phytopathogen (92,93). In particular, the acquisition and exchange of CWDEs and XLPs might underlie the double emergence of phytopathogenicity in *Arsenophonus*. Biochemical characterization of these enzymes and their substrates will be required to further support this hypothesis.

## Materials and methods

### DNA extraction for insect metagenomes

For Pf-FR, adult *C. wagneri* specimens were collected in a strawberry field in Dordogne (France) in June 2019. For Ap-FR, adult *P. leporinus* specimens were collected during the first SBR outbreak in Burgundy (France) in the early 2000s. For Ap-CH, adult *P. leporinus* specimens were collected in a sugar beet field in Gilly (Switzerland) in 2020. Insects from France were surface-sterilized by serial washes in 20% bleach, sterile water, 70% ethanol and sterile water. Individual insects were then ground in 400 µl 2% CTAB buffer (2% CTAB, 2% PVP K40, 1.4 M NaCl, 20 mM EDTA, 100 mM Tris-HCl, 0.02% β-mercaptoethanol) and incubated at 65°C for 1.5 hours. The DNA was then extracted as previously described (58). No surface-sterilization was performed for insects collected in Switzerland, and they were stored in 70% ethanol at -20°C until further use. For these specimens, DNA was extracted using a 3% CTAB protocol (14). All DNA samples were treated with 20 µg RNase A at 37°C for 30 min, followed by chloroform/isoamyl alcohol extraction and isopropanol precipitation. DNA pellets were resuspended in 40 µl of 1X TE buffer for samples from France, whereas 100 µl of DNase-free water was used for Swiss samples.

### Quantification of bacterial titre

Bacterial titres were estimated in DNA samples by qPCR. For Pf-FR and Ap-FR, samples were normalized to 50 ng/µl to compare Ct values between samples. All samples were tested in duplicates with the previously published primers and FAM-labelled TaqMan probes targeting the *spoT* gene of Pf (4). For Ap-FR, primers and probe (SBR-F/R/FAM) are listed in Table S1. Reactions were performed as previously described (58). For samples from Switzerland, a similar qPCR was performed using a different protocol (14). Samples with the highest bacterial titres were chosen for high-throughput sequencing.

### Genome sequencing and assembly

For Ap-CH and Pf-FR, long-read sequencing libraries were prepared using the Ligation Sequencing kits SQK-LSK 110 and 109 (Oxford Nanopore Technologies, UK), respectively. Each library was sequenced on an R9.4 flowcell on the MinION sequencer, producing 16 and 6 Gb of data, respectively. Basecalling was done using Guppy v5.0.11 (in high-accuracy mode) and only ≥ 500 bp-reads passing the Q7 quality filter were retained. In addition, 2 x 150 bp paired-end reads were obtained from an Illumina Novaseq platform (Macrogen), producing 50 million and 476 million reads for Ap-CH and Pf-FR, respectively. The reads were quality-trimmed using Trimmomatic v0.38 (5), retaining only reads ≥ Q30.

Reads belonging to *Arsenophonus* were extracted from the datasets via mapping (using Minimap2 v2.15 (6) for the long reads and bwa mem v0.7.17 for the short reads, respectively) against a database of all published *Arsenophonus* genomes (as of September 2023, Table 1). Several hybrid assemblers were then tested: Masurca v4.0.7 (7, 8), Spades v3.15.1 (9) and Unicycler v0.4.9 (10). For Pf-FR, the most contiguous assembly was obtained using Masurca, and was further polished with short reads using Polca (part of the Masurca package) and scaffolded using two iterations of SSPACE v2.1.1 (11). For Ap-CH, the most contiguous assembly was obtained using Unicycler, and this assembly was scaffolded using two iterations of SSPACE.

The genome of Ap-FR was assembled only from short reads since the samples had been stored for more than 15 years prior DNA extraction, resulting in highly fragmented DNA precluding long-read sequencing. Illumina sequencing was performed as mentioned above, producing 486 million 150 bp paired-end reads. Following quality trimming, the reads were assembled using Megahit v1.2.9 (12) and contigs belonging to Ap were identified using BlobTools v1.1.1 (https://github.com/DRL/blobtools) and mapping against a database of all published *Arsenophonus* genomes using bwa mem. The contigs were scaffolded using Redundans (13) using the Ap-CH assembly as reference, followed by one iteration of SSPACE and Gapfiller v2.1.2 (14). Coverages were determined with Mosdepth v0.3.4 (94). CheckM v1.1.6 (95) was used to assess genome completeness. Synteny blocks were identified with Sibelia v3.0.7 (96).

### Functional annotation

The three assemblies were annotated using the NCBI PGAP v2023-05-17.build6771. COG categories and KEGG Ontology terms were determined using eggNOG-mapper v2 (15). Phage regions were detected with Phaster (97). Schematic representations of genetic modules were obtained with Clinker (98). Circular representations of the scaffolds were produced with the R package Circlize v0.4.10 (99). Heatmaps were constructed using the ComplexHeatmap R package (100). Biosynthetic capacities were evaluated using KeggDecoder v1.3 (101), TXSscan v1.1.0 (102) and antiSMASH v7.beta (103). Toxins and effectors were detected by Blastp searches in a local database of *Arsenophonus* proteomes.

### Phylogenomics and taxonomy

Orthofinder v2.5.4 (16) was used to identify single-copy orthologs shared between Pf-FR, Ap-CH, Ap–FR and 22 *Arsenophonus* genomes published as of September 2023 (Table 1). *Proteus mirabilis* HI4320 (GCF_000069965.1) and *Providencia stuartii* MRSN 2154 (GCF_000259175.1) were used as outgroups. Amino acid sequences of each gene were aligned using Muscle v3.1.31 (17), followed by concatenation with geneStitcher.py (https://github.com/ballesterus/Utensils/blob/master/geneStitcher.py). IQ-TREE v1.6.12 (18) was used to predict the best substitution model for each partition (19, 20) and to produce a ML phylogenetic tree with 1000 bootstrap iterations. The tree was edited in iTOL (104). ANI values were obtained using the enveomics toolbox collection (105).

### Pangenome analysis

An orthology clustering analysis was conducted using Orthofinder v2.5.4 (106) on all *Arsenophonus* genomes used for the phylogenomic analyses. Shared OGs were identified using the R package UpSetR v1.4.0 (107). To further analyse the genes present only in Ap and/or Pf, Blastp searches were conducted on the longest sequence for each group. These searches excluded additional undetected pseudogenes and orthologs shared with recently-published *Arsenophonus* genomes reconstructed from SRA datasets and endosymbionts of louse flies (44,108). For OG shared by Ap and Pf, functional domains were identified with MotifFinder (https://www.genome.jp/tools/motif/). *In silico* predictions were conducted using SignalP v6.0 (109), SecretomeP v2.0 (110), PSORTb v3.0.3 (111). Phylogenetic analyses were performed as previously described (112).

### Gene expression analyses

RT-qPCR analyses were conducted for several Ap-CH genes in *P. leporinus* females and periwinkle stems. Insects were collected at the adult stage, after a rearing period of *c.* 5 months in controlled conditions (113). Periwinkle seedlings were inoculated with field-collected insects as previously described (14), then maintained in insect-proof cages in greenhouse conditions for three months. Whole insect bodies and plant stem samples were freeze-dried in liquid nitrogen and stored at – 80°C until further use. Total RNA was extracted using a 3% CTAB protocol (14), treated with DNase and quantified using a Qubit fluorometer. cDNA was synthetized for 1 μg of each RNA sample using random hexamers (Invitrogen) in combination with the Superscript reverse transcriptase (Promega). qPCR assays were conducted as previously described (14), using primers and probes listed in Table S1. Expressions were obtained for five biological replicates and were normalized using the Ap-CH *gyrase* or *rpoB* genes. qPCR efficiencies were assessed based on five to six ten-fold serial dilutions of an infected sample in healthy RNA extracts. Specificity was checked by gel electrophoresis.

## Author contributions

MM, CD, JB, DR, XF and JD prepared samples and conducted HTS analyses. MM conducted the gene expression analyses. MM and JD performed the bioinformatics analyses and wrote the initial draft. CD, DR and RG provided technical assistance. JD, FF, XF and OS obtained funding. All authors reviewed the manuscript and agreed to publication.

## Acknowledgements

We would like to thank Floriane Bussereau (Agroscope, Switzerland) and Frédéric Gatineau (Cirad, France) for providing insects. This research was funded by the European Union’s Horizon 2020 research and innovation programme under the Marie Sklodowska-Curie (grant agreement No 792813 to JD) and the Swiss Federal Office for Agriculture (Grant 2020/33/LES-Z II to OS).

